## Supplementary material for "Uneven terrain treadmill walking in younger and older adults": S1 Fig. Obstacle positioning on treadmill.

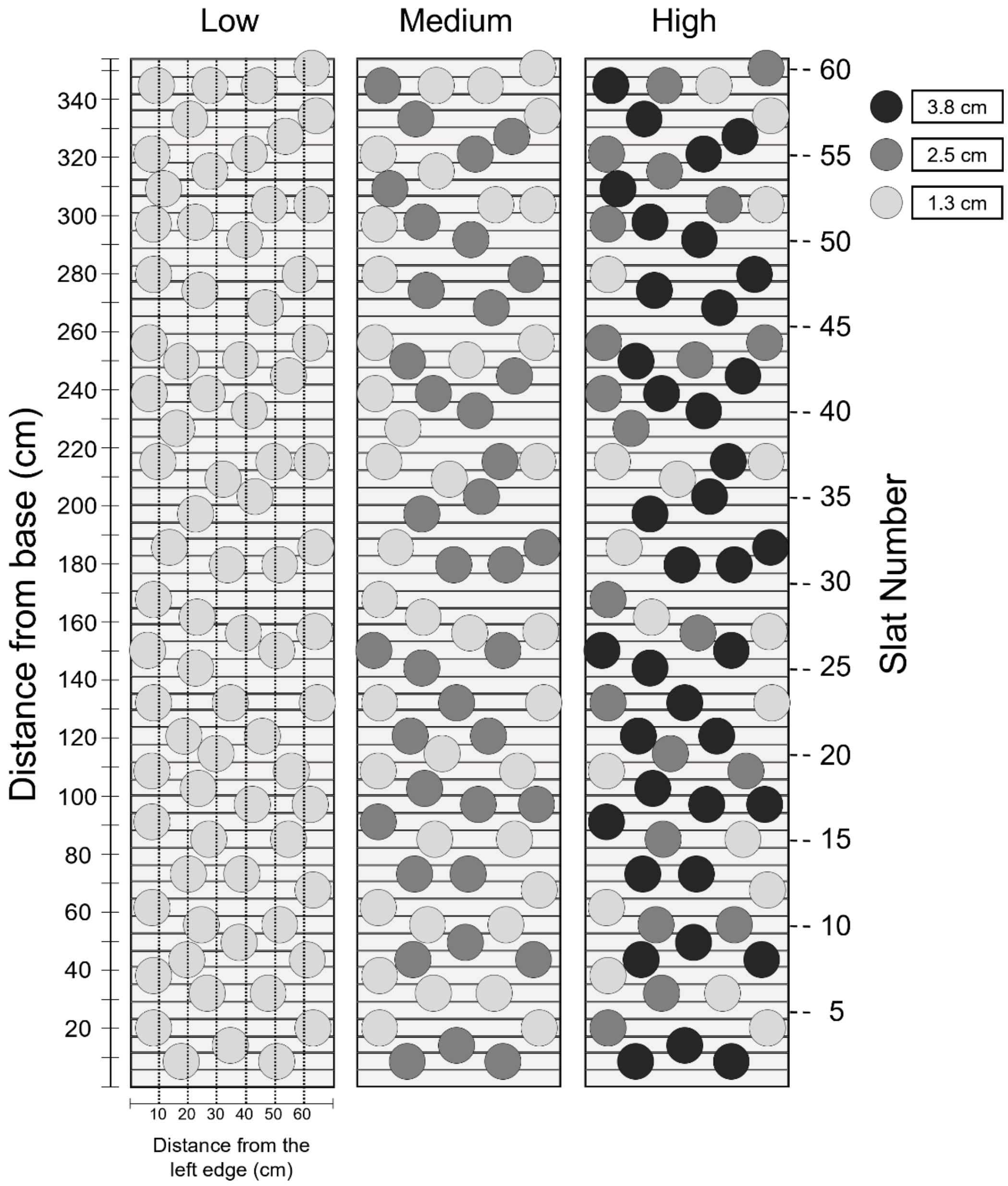


**S1 Fig. Obstacle positioning on treadmill.** Obstacles were placed systematically so that the terrain conditions were repeatable across participants. The total weight of the added foam obstacles was 3.5, 5.2, and 7.9 pounds (1.6, 2.4, 3.6 kg) for the Low, Medium, and High conditions, respectively.
