## Supplementary material for "Uneven terrain treadmill walking in younger and older adults": S2 Table. Statistical model results for step duration.

**S2 Table. Statistical model results for step duration (s) after accounting for walking speed.**

| | Value | Std. Error | DF | t-value | p-value | Sig. | ES |
|---|---|---|---|---|---|---|---|
| **Intercept** | | | | | | | 0.14 |
| HFOA, Flat | 0.131 | 0.0554 | 200 | 2.36 | 0.0192 | * | |
| **Group** | | | | | | | 0.46 |
| YA | -0.259 | 0.0871 | 200 | -2.97 | 0.0033 | * | |
| LFOA | 0.369 | 0.0756 | 200 | 4.89 | 0.0000 | * | |
| **Terrain** | | | | | | | 0.11 |
| Low | -0.079 | 0.0783 | 200 | -1.01 | 0.3157 | | |
| Medium | -0.100 | 0.0783 | 200 | -1.28 | 0.2033 | | |
| High | -0.142 | 0.0783 | 200 | -3.37 | 0.0706 | | |
| **Interaction** | | | | | | | 0.06 |
| YA Low | 0.089 | 0.1231 | 200 | 0.73 | 0.4688 | | |
| YA Medium | 0.096 | 0.1231 | 200 | 0.78 | 0.4371 | | |
| YA High | 0.111 | 0.1231 | 200 | 0.90 | 0.3675 | | |
| LFOA Low | 0.020 | 0.1074 | 200 | 0.18 | 0.8558 | | |
| LFOA Medium | 0.037 | 0.1069 | 200 | 0.35 | 0.7280 | | |
| LFOA High | 0.069 | 0.1074 | 200 | 0.64 | 0.5201 | | |

DF, degrees of freedom; ES, Effect Size; HFOA, higher-functioning old adults; LFOA = lower-functioning old adults; YA, young adults.
