## Supplementary material for "Uneven terrain treadmill walking in younger and older adults": S3 Table. Statistical model results for step duration variability.

**S3 Table. Statistical model results for step duration variability (%) after accounting for walking speed.**

| | Value | Std. Error | DF | t-value | p-value | Sig. | ES |
|---|---|---|---|---|---|---|---|
| **Intercept** | | | | | | | 0.11 |
| HFOA, Flat | -1.41 | 0.747 | 200 | -1.89 | 0.0600 | | |
| **Group** | | | | | | | 0.46 |
| YA | -3.36 | 1.175 | 200 | -2.86 | 0.0047 | * | |
| LFOA | 5.06 | 1.020 | 200 | 4.96 | 0.0000 | * | |
| **Terrain** | | | | | | | 0.22 |
| Low | 1.66 | 1.057 | 200 | 1.57 | 0.1182 | | |
| Medium | 2.16 | 1.057 | 200 | 2.04 | 0.0422 | * | |
| High | 3.86 | 1.057 | 200 | 3.65 | 0.0003 | * | |
| **Interaction** | | | | | | | 0.15 |
| YA Low | 0.17 | 1.661 | 200 | 0.10 | 0.9204 | | |
| YA Medium | 0.97 | 1.661 | 200 | 0.59 | 0.5583 | | |
| YA High | 0.33 | 1.661 | 200 | 0.20 | 0.8411 | | |
| LFOA Low | 0.96 | 1.448 | 200 | 0.66 | 0.5071 | | |
| LFOA Medium | 1.79 | 1.442 | 200 | 1.24 | 0.2151 | | |
| LFOA High | 3.20 | 1.448 | 200 | 2.21 | 0.0284 | * | |

DF, degrees of freedom; ES, Effect Size; HFOA, higher-functioning old adults; LFOA = lower-functioning old adults; YA, young adults.
