## Supplementary material for "Uneven terrain treadmill walking in younger and older adults": S4 Table. Statistical model results for anteroposterior excursion variability.

**S4 Table. Statistical model results for anteroposterior excursion variability (%) after accounting for walking speed.**

| | Value | Std. Error | DF | t-value | p-value | Sig. | ES |
|---|---|---|---|---|---|---|---|
| **Intercept** | | | | | | | 0.02 |
| HFOA, Flat | -0.62 | 1.84 | 200 | -0.34 | 0.7327 | | |
| **Group** | | | | | | | 0.64 |
| YA | -13.45 | 2.89 | 200 | -4.66 | 0.0000 | * | |
| LFOA | 16.21 | 2.51 | 200 | 6.46 | 0.0000 | * | |
| **Terrain** | | | | | | | 0.21 |
| Low | 3.66 | 2.60 | 200 | 1.41 | 0.1603 | | |
| Medium | 5.15 | 2.60 | 200 | 2.04 | 0.0422 | * | |
| High | 9.03 | 2.60 | 200 | 3.47 | 0.0006 | * | |
| **Interaction** | | | | | | | 0.04 |
| YA Low | -0.81 | 4.09 | 200 | -0.20 | 0.8421 | | |
| YA Medium | -0.17 | 4.09 | 200 | -0.04 | 0.9664 | | |
| YA High | -2.10 | 4.09 | 200 | -0.52 | 0.6071 | | |
| LFOA Low | 0.21 | 3.56 | 200 | 0.06 | 0.9522 | | |
| LFOA Medium | 0.02 | 3.55 | 200 | 0.01 | 0.9949 | | |
| LFOA High | -1.60 | 3.56 | 200 | -0.45 | 0.6531 | | |

DF, degrees of freedom; ES, Effect Size; HFOA, higher-functioning old adults; LFOA = lower-functioning old adults; YA, young adults.
