## Supplementary material for "Uneven terrain treadmill walking in younger and older adults": S5 Table. Statistical model results for mediolateral excursion variability.

**S5 Table. Statistical model results for mediolateral excursion variability (%) after accounting for walking speed.**

| | Value | Std. Error | DF | t-value | p-value | Sig. | ES |
|---|---|---|---|---|---|---|---|
| **Intercept** | | | | | | | 0.14 |
| HFOA, Flat | -2.84 | 1.22 | 200 | -2.33 | 0.0210 | * | |
| **Group** | | | | | | | 0.26 |
| YA | -1.98 | 1.92 | 200 | -1.03 | 0.3038 | | |
| LFOA | 5.57 | 1.67 | 200 | 3.34 | 0.0010 | * | |
| **Terrain** | | | | | | | 0.27 |
| Low | 2.76 | 1.73 | 200 | 1.59 | 0.1127 | | |
| Medium | 5.55 | 1.73 | 200 | 3.21 | 0.0016 | * | |
| High | 7.23 | 1.73 | 200 | 4.18 | 0.0000 | * | |
| **Interaction** | | | | | | | 0.08 |
| YA Low | -0.02 | 2.72 | 200 | 0.01 | 0.9928 | | |
| YA Medium | -2.36 | 2.72 | 200 | -0.87 | 0.3855 | | |
| YA High | -2.45 | 2.72 | 200 | -0.90 | 0.3682 | | |
| LFOA Low | 1.02 | 2.37 | 200 | 0.43 | 0.6670 | | |
| LFOA Medium | -0.95 | 2.36 | 200 | -0.40 | 0.6881 | | |
| LFOA High | -0.58 | 2.37 | 200 | -0.24 | 0.8070 | | |

DF, degrees of freedom; ES, Effect Size; HFOA, higher-functioning old adults; LFOA = lower-functioning old adults; YA, young adults.
