## Supplementary material for "Uneven terrain treadmill walking in younger and older adults": S6 Table. Statistical model results for perceived stability ratings.

| | OR | 2.5% | 97.5% | Sig. |
|---|---|---|---|---|
| **Group** | | | | |
| YA | 1.22 | 0.721 | 2.05 | |
| LO | 2.83 | 1.798 | 4.50 | * |
| **Terrain** | | | | |
| Low | 4.15 | 1.799 | 10.82 | * |
| Medium | 16.96 | 7.830 | 42.56 | * |
| High | 68.03 | 31.010 | 172.67 | * |

HO, higher-functioning old adults; LO = lower-functioning old adults; OR, odds ratio; YA, younger adults.
If odds ratio > 1, the effect is in the direction of less stable walking.
If odds ratio < 1, the effect is in the direction of more stable walking.
If odds ratio confidence interval [2.5%, 97.5%] contains 1, the effect is not significant.
